# Agentic transcranial functional ultrasound imaging: @fUS

**DOI:** 10.64898/2026.09.25.752961

**Authors:** Zihao Chen, Zexin Yuan, Zhaoqi Lu, Na Li, Ceyi Fu, Le Sun, Hongying Yao, Huizhu Liu, Quanxiang Xian, Tianyi Wang, Wei Li, Zirui Liang, Boxing Li, Ying Zhang, Ying Li, Hongsheng Wang, Wen-Jie Bian, Xiaodong Liu, Yan Chen, Bo Li, Lei Sun, Yimin Wang, Zhihai Qiu

## Abstract

Functional ultrasound (fUS) imaging provides deep, wide-field access to brain function, but instrument cost and complex acquisition and analysis workflows restrict its wider adoption. Here we introduce @fUS, an agentic transcranial imaging platform with an open architecture that integrates purpose-built hardware with executable experimental skills and persistent structured memory. The system combines 128-channel, 16-bit acquisition at 125 MHz with GPU-based processing and enables functional imaging through the intact scalp and skull of mice, with approximately 100-μm spatial resolution and 10-Hz temporal sampling, providing a basis for longitudinal studies without cranial-window surgery. Its agent connects scientific objectives to experimental design, direct instrument control and data analysis, retaining context across interactions and supporting inspectable workflows through text and hands-free voice control. Neuroscience trainees with limited engineering experience independently configured the system within 1 hour. Agent-assisted exploration of whisker-stimulation datasets revealed low-frequency vascular changes beyond conventional response mapping, supported by complementary two-photon measurements of single-vessel diameter. By combining transcranial imaging with accessible instrument control and analysis, @fUS provides a foundation for wider adoption and larger, more diverse neuroscience datasets. More broadly, it offers a framework for scientific instruments in which measurement, computation and experimental reasoning are developed as parts of the same system.

## Introduction

Scientific instruments have transformed our ability to observe nature, yet translating scientific questions into measurements still relies heavily on human expertise^1^. Connecting computational reasoning with experimental hardware is a key next step for artificial intelligence (AI) in science^2–5^. This connection could enable a continuous cycle in which scientific objectives guide physical measurements and the resulting evidence informs subsequent actions^6–8^. Realizing this vision requires instruments whose capabilities, data and control interfaces are accessible to both researchers and intelligent agents, making openness and programmability central to instrument design^9–11^.

Functional ultrasound (fUS) imaging illustrates the need for such integration. By measuring cerebral hemodynamics associated with neural activity, fUS provides sensitive functional measurements across wide fields of view and deep brain structures^12–16^. Its combination of spatial resolution, temporal sampling and imaging depth complements magnetic resonance and optical methods, supporting systems and developmental neuroscience^17–22^. However, wider adoption remains constrained by instrument cost and technical complexity^23–26^. Commercial systems can restrict access to raw data and low-level processing, whereas programmable ultrasound platforms require substantial adaptation and expertise in electronics, signal processing and software development to meet the throughput demands of functional imaging^16,23,24^.

The challenge extends beyond access to hardware and code. fUS experiments require coordinated choices about acoustic parameters, acquisition sequences, beamforming, clutter rejection, quality control and statistical analysis^23,25,27–31^. Much of the knowledge needed to make these choices, recognize artifacts and troubleshoot failures is acquired through experience and remains difficult to transfer between laboratories^23,25,30,32^. Open hardware and software therefore do not, by themselves, make an experimental capability broadly accessible^33,34^. Scaling fUS requires ways to share practical expertise together with the instrument.

Agentic AI offers a way to make such expertise operational. Large language models can combine heterogeneous information and invoke external tools, while agent architectures organize multistep actions and retain context^3,5,35–38^. Recent systems have connected agents to bioimage analysis and microscope control through software interfaces^39,40^. For complex instruments such as fUS, this approach can extend to the design of the hardware control and acquisition architecture itself, exposing acquisition settings, instrument state, raw signals and reconstruction operations through programmable interfaces. Agents could then inspect intermediate measurements and use analytical outcomes to inform subsequent acquisitions^4^. Reconstructed images could also be linked to specialist analysis environments such as CopilotJ^41^.

Here we introduce @fUS, an open, agentic transcranial imaging platform that integrates purpose-built fUS hardware with programmable acquisition, reconstruction and analysis. The system enables functional imaging through the intact mouse skull without skull replacement, providing a less invasive route to repeated measurements and longitudinal studies. Neuroscience trainees with limited engineering experience independently configured the system within 1 hour. Its agent combines executable experimental skills with persistent structured memory to support hypothesis generation, experimental design, instrument operation, quality control and data exploration across sessions. Voice interaction enables hands-free control during experimental procedures. Agent-assisted exploration of whisker-stimulation datasets revealed low-frequency vascular changes supported by complementary two-photon measurements. This integration provides a foundation for broader access to fUS and for scientific instruments that connect reasoning directly to experimental execution.

## Results

### An agentic fUS platform

@fUS was developed to connect researcher-defined questions with ultrasound acquisition, reconstruction and analysis (Fig. 1). Purpose-built imaging hardware was integrated with an agent through which instrument-control and analytical tools could be invoked. RF signals were digitized by the 128-channel electronics at 16 bits and 125 MHz and transferred to a workstation for GPU reconstruction, with a measured maximum transfer rate of 16 GB/s to host memory. Raw RF data were retained so that measurements could be revisited through alternative reconstructions. Hardware components and the control architecture are detailed in Fig. S1A, B and Methods.

**Fig. 1.**
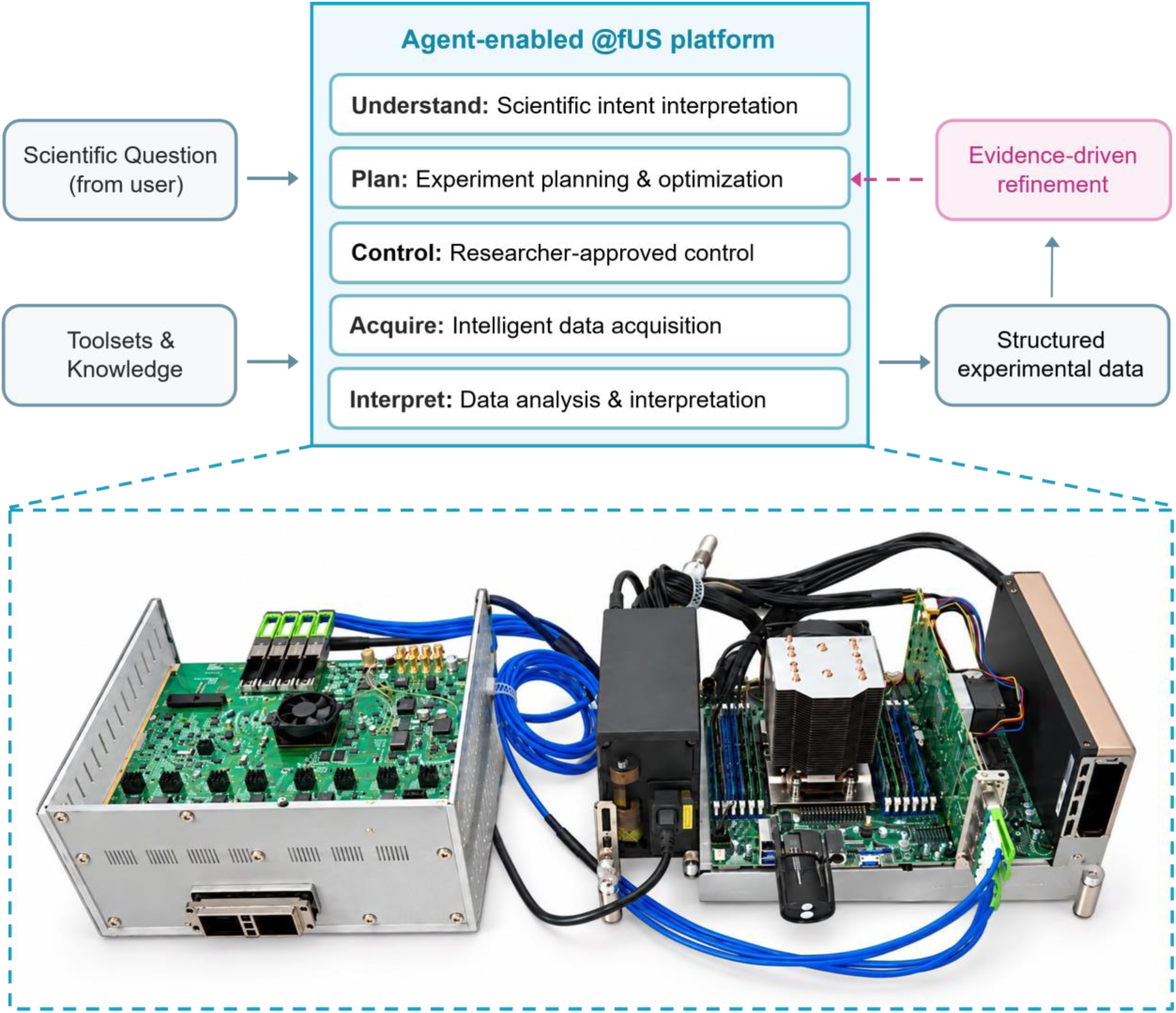
AI reasoning is connected to physical experimentation through @fUS. Scientific questions are linked to experimental planning, instrument control, acquisition and analysis through the @fUS platform. Instrument tools are combined with locally stored knowledge and experimental context by the agent, and proposed operations are submitted for researcher approval. Configuration and timing metadata are retained with recordings, so that subsequent questions and settings can be informed by analytical findings. The custom ultrasound electronics and acquisition and processing workstation are shown in the photograph. Hardware and agent implementation are detailed in Fig. S1.

Researcher requests were linked to instrument and analytical tools through executable skills, and acquisition and reconstruction settings were exposed through text-based configurations. Experimental context was retained across interactions in locally stored documentation, datasets and persistent memory. Recordings were linked to acquisition conditions and subsequent analyses through configuration and timing metadata. Proposed operations could be inspected, approved and revised by researchers through text and voice interfaces; manual control was also supported. Experimental planning was thereby connected with data interpretation (Supplementary Video 1-4; Fig. S1B, C and Table S1). In practice, @fUS was independently installed, configured and operated within 1 h by neuroscience trainees with limited engineering experience (Supplementary Video 6-7; Fig. S1D).

The agent-assisted workflow was evaluated through parameter comparisons, transcranial functional imaging and exploration of vascular dynamics. Acquisition and reconstruction settings were first examined for their effects on vascular image quality and resource demand. This configuration workflow was then applied to transcranial imaging before agent assistance was extended to the exploration of experimental findings.

### Agent-guided imaging configuration control

In fUS, vascular visibility, temporal sampling and computational demand are jointly determined by acquisition and reconstruction settings. Direct access to these choices is provided to the agent through structured configuration files, instrument-state readouts and image-quality outputs (Fig. 2A and Table S1). Imaging objectives and resource constraints are translated by the agent into candidate TXT and CSV configurations; images and quantitative comparisons are then organized after researcher-approved execution. Parameter selection is supported by weighing vascular visibility and contrast against memory, processing and temporal-sampling constraints.

**Fig. 2.**
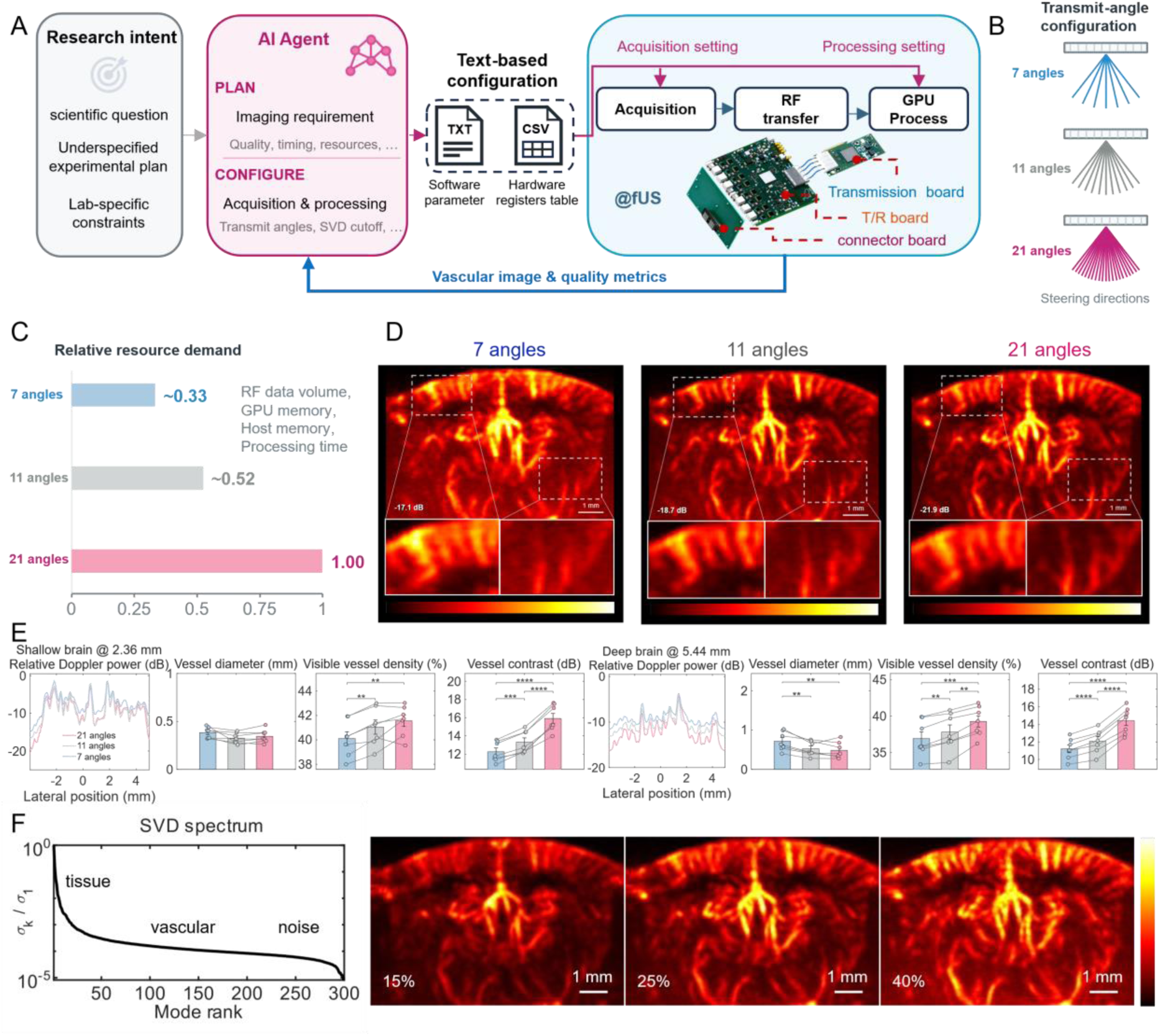
Agent-guided configuration balances vascular image quality with resource demand and spatiotemporal imaging requirements. A, Workflow from research intent to imaging configuration and feedback. The scientific question, initial plan and laboratory constraints are translated by the agent into proposed imaging requirements and acquisition and processing settings. Control over acquisition and GPU reconstruction is exposed through TXT software parameters and CSV hardware-register tables. Following validation and researcher approval, acquisition, RF transfer and processing are executed by the host. Further review and approved changes are guided by vascular images, quality metrics and resource constraints. B, Transmit-angle configurations used for separate acquisitions: seven angles spanning ±6°, eleven spanning ±10° or twenty-one spanning ±10°. C, Schematic angle-dependent resource demand, normalized to 21 angles. Measured system resource use and real-time Doppler imaging frame rates are reported in Fig. S2. D, Representative power Doppler images, with enlarged superficial and deep regions. E, Lateral power profiles at depths of 2.36 and 5.44 mm, apparent vessel diameter (−3-dB profile width), visible vessel area fraction and vessel-to-background power contrast. Mean ± s.e.m. is shown by bars; individual mice are represented by connected points (n = 7; ≥100 images per mouse and configuration, averaged within each mouse). Two-sided paired t-tests were performed with Holm correction across three angle pairs within each region and metric; **P < 0.01, ***P < 0.001 and ****P < 0.0001. F, Normalized SVD spectrum and reconstructions after removal of the leading 15%, 25% or 40% of components by rank. Conceptual tissue, vascular and noise contributions are indicated by spectrum labels. Seven, eleven and twenty-one angles are denoted by blue, grey and pink. Imaging was performed through cranial windows under isoflurane anesthesia. A complementary retrospective comparison of spatial and temporal measurements is shown in Fig. S3. GPU, graphics processing unit; T/R, transmit–receive. Scale bars, 1 mm (D, F).

Twenty-one-angle acquisition was supported by the streaming architecture, with a measured maximum transfer rate of 16 GB/s to host memory. Separate recordings were acquired with 7, 11 and 21 transmit angles and reconstructed using 300-frame ensembles (Fig. 2B; Methods). Angular ranges of ±6° and ±10° were covered by the seven-angle acquisition and the other configurations, respectively. Apparent vessel diameter, visible vessel area fraction and vessel-to-background power contrast were quantified by the agent. Relative resource demand was estimated from angle count and normalized to the 21-angle configuration (Fig. 2C). In resource tests at a fixed pulse repetition frequency, host RAM and GPU memory usage were reduced from 41.08 to 17.15 GiB and from 38.79 to 13.48 GiB, respectively, between 21 and 7 angles (Fig. S2). Finer temporal sampling was obtained with fewer angles: real-time Doppler imaging frame rates of 1.82, 3.47 and 5.45 Hz were measured for 21, 11 and 7 angles, respectively. Shorter RF preparation times were also measured, whereas higher CPU and GPU utilization and GPU power consumption were observed. Under the experimental conditions and resource constraints, image-quality and resource measurements were compared by the agent to weigh vascular signal discrimination against temporal resolution and support researcher-approved parameter selection.

Vascular branches were delineated more clearly in higher-angle reconstructions, with fine vessels resolved against the background (Fig. 2D). Across seven mice, superficial apparent vessel diameters were 0.384 ± 0.022, 0.325 ± 0.021 and 0.345 ± 0.026 mm for 7, 11 and 21 angles, respectively (mean ± s.e.m.; Fig. 2E). None of the pairwise differences were significant after Holm correction (adjusted P = 0.0886 for 7 versus 11 angles and 0.182 for both comparisons involving 21 angles). Deep-vessel diameters were 0.717 ± 0.091, 0.525 ± 0.069 and 0.485 ± 0.078 mm, respectively. Narrower profiles were obtained with both 11- and 21-angle reconstructions than with seven-angle reconstruction (adjusted P = 0.00453 and 0.00357), with no significant difference between 11 and 21 angles (adjusted P = 0.373). Angular effects on apparent width were therefore most evident in deep-vessel profiles reconstructed with seven angles.

Visible vessel area fractions were 40.11 ± 0.56%, 41.04 ± 0.58% and 41.56 ± 0.47% superficially, and 36.93 ± 0.93%, 37.82 ± 0.98% and 39.28 ± 0.78% in the deep region for 7, 11 and 21 angles, respectively. Greater superficial coverage was measured with both 11 and 21 angles relative to seven angles (adjusted P = 0.00154 and 0.00155), while no significant difference was detected between 11 and 21 angles (adjusted P = 0.168). Greater deep-region coverage was measured between successive configurations (adjusted P = 0.00240 for both 7 versus 11 and 11 versus 21 angles), with adjusted P = 2.49 × 10⁻⁴ for 7 versus 21 angles.

Progressively greater vessel-to-background power contrast was measured across the three configurations. For 7, 11 and 21 angles, superficial contrast was 12.26 ± 0.41, 13.32 ± 0.47 and 15.92 ± 0.55 dB, and deep-region contrast was 11.20 ± 0.44, 12.07 ± 0.50 and 14.42 ± 0.57 dB, respectively. All pairwise comparisons were significant: superficial adjusted P values were 1.13 × 10⁻⁴ for 7 versus 11 angles, 1.95 × 10⁻⁵ for 11 versus 21 angles and 1.47 × 10⁻⁵ for 7 versus 21 angles; corresponding deep-region values were 7.15 × 10⁻⁵, 1.07 × 10⁻⁵ and 1.07 × 10⁻⁵. Comparisons were performed using two-sided paired t-tests on mouse-level summaries, with Holm correction within each region and metric. The gains in visible vessel area and contrast were consistent with improved delineation of weak vascular signals, including small vessels in deeper regions (Fig. 2D, E).

Clutter rejection was examined as a further selection dimension. Fine-vessel visibility and background intensity were altered by removal of the leading 15%, 25% or 40% of singular value decomposition (SVD) components by rank (Fig. 2F). Some fine vessels were less clearly resolved at the lowest rejection fraction, whereas stronger deep-region background noise was observed at the highest fraction. Agent-assisted selection of angular compounding and SVD rejection was thus informed by image-quality comparisons and measured resource demands within researcher-defined memory, processing and temporal-sampling constraints.

To reduce confounding by changes in mouse physiological state between separate acquisitions, a complementary comparison was performed by retrospectively reconstructing 7-, 11- and 21-angle images from the same RF recordings (Fig. S3). Greater intensity variation with increasing angle count was detected in the agent-executed temporal analyses (Fig. S3A–E). In a representative 190-frame sequence, mean inter-frame differences of 13.62%, 14.59% and 16.86% were measured for 7, 11 and 21 angles, respectively. Mean adjacent-frame spatial correlations of 0.9746, 0.9748 and 0.9779 were measured in the same order (Fig. S3E), while central-section position jitter of 93.45, 71.50 and 58.77 µm was measured in a representative cortical vessel (Fig. S3I, K). The combination of greater intensity variation and more consistent spatial structure was considered compatible with retention of vascular fluctuations carrying additional functional information. Recovery of localized, stimulus-associated responses through the intact scalp and skull was examined next.

### Agent-assisted transcranial functional imaging

This configuration workflow was next applied to imaging through the intact mouse scalp and skull (Fig. 3). Reconstruction comparisons and vascular and functional quantification were assisted by the agent under researcher supervision.

**Fig. 3.**
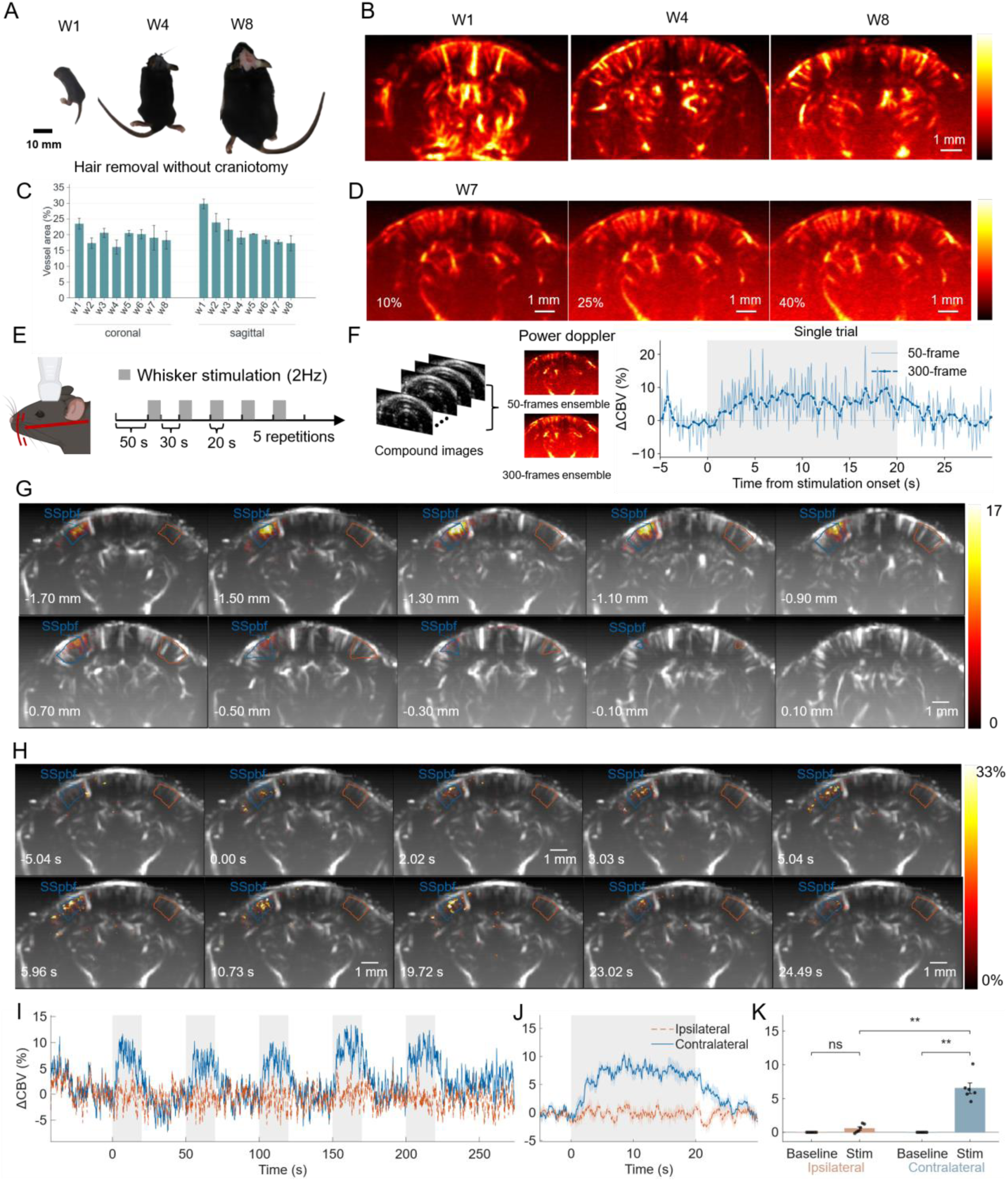
Agent-assisted fUS maps cerebral vessels and evoked responses through the intact scalp and skull. A, B, Intact-scalp and intact-skull preparations (A) and representative power Doppler images at W1, W4 and W8 (B). Hair was removed only after 2 weeks of age. C, Coronal and sagittal visible vessel area fractions across W1–W8 (mean ± standard deviation; n = 3 mice per age group). The leading 20% of SVD components were rejected by rank in B, C. D, W7 reconstructions with 10%, 25% or 40% rank rejection. Normalized power Doppler signal is encoded by colors in B, D. E, A 50-s baseline followed by five blocks of 20-s whisker stimulation at 2 Hz and 30-s recovery. F, Reconstructions and unsmoothed responses from the same single trial using 50- or 300-frame ensembles of identical RF data, sampled at intervals of 0.0917 or 0.5503 s (approximately 10.9 or 1.82 Hz), respectively. The trade-off between temporal sampling and baseline variability is illustrated at fixed acquisition settings. G, GLM-derived statistical Z maps across ten coronal positions at 200-µm intervals, from −1.70 to +0.10 mm relative to bregma. H, Five-trial-averaged, nuisance-adjusted ΔCBV within the full-session GLM significance mask, displayed on a common 0–33% scale. GLM maps were generated using one-sided tests with Benjamini–Hochberg correction within each slice (q < 0.05). Contralateral and ipsilateral SSp-bf are outlined in blue and orange in G, H. I, J, Continuous (I) and stimulus-aligned (J) responses from the plane shown in H; mean ± s.e.m. across five trials is shown by lines and shading in J. Stimulation is indicated by grey shading. K, Six-mouse responses using each animal’s plane with the largest five-trial mean contralateral response (bars, mean ± s.e.m.; points, mice). Two-sided paired t-tests with Holm correction: ipsilateral baseline versus stimulation, P = 0.0837; contralateral baseline versus stimulation and contralateral versus ipsilateral responses, both P = 0.00110. ns, not significant; **P < 0.01. This response-selected analysis is exploratory. Vascular imaging was performed under isoflurane; functional recordings were acquired in W7 mice under Zoletil–xylazine anesthesia. Analyses were performed by the agent using researcher-approved MATLAB workflows (Methods). Stimulation onset is denoted by time zero. Scale bars, 10 mm (A) and 1 mm (B, D, G, H).

With the leading 20% of SVD components removed by rank, intracranial vessels were resolved by transcranial power Doppler imaging at 1, 4 and 8 weeks of age without craniotomy or skull replacement (Fig. 3A, B). Visible vascular coverage was quantified in coronal and sagittal images across W1–W8 (n = 3 mice per weekly age group; Fig. 3C). Hair was removed only in mice older than 2 weeks.

In a representative W7 mouse, vessel visibility and background signal were altered by rejection of the leading 10%, 25% or 40% of SVD components (Fig. 3D). Clutter-filtering choices could therefore be examined through the agent-accessible reconstruction workflow.

Functional imaging was evaluated during five 20-s blocks of 2-Hz whisker stimulation in 7-week-old mice under Zoletil–xylazine anesthesia to facilitate immobilization (Fig. 3E; Methods). Sampling intervals of 0.0917 and 0.5503 s, corresponding to approximately 10.9 and 1.82 Hz, were obtained by reconstruction of the same RF data with 50- and 300-frame ensembles (Fig. 3F). In the displayed single trial, stimulation-period mean ΔCBV values of 5.76% and 5.71% were measured, respectively, with baseline temporal standard deviation values of 4.27 and 2.37 percentage points. Baseline variability was thus reduced with the longer ensemble, while the sampling interval was increased sixfold. The 50-frame reconstruction was then evaluated for localized, repeatable stimulus-associated responses.

Images were aligned to recorded triggers; responses were extracted from anatomical ROIs and general linear model (GLM) activation maps were generated by the agent. Across ten coronal planes spaced 200 µm apart, from −1.70 to +0.10 mm relative to bregma, activation was localized predominantly to the contralateral primary somatosensory barrel field (SSp-bf; Fig. 3G). Differences in response location and extent were observed across planes with different cross-sectional anatomy of SSp-bf. At −1.50 mm in the representative mouse, significant positive activation was detected over 0.815 mm², or 82.29% of contralateral SSp-bf. Anatomical localization of the functional response through the intact preparation was supported by these spatially varying maps.

The temporal evolution of the response was resolved in stimulus-aligned maps (Fig. 3H). Nuisance-adjusted means across contralateral SSp-bf of 4.00% at 2.02 s, 9.66% at 5.96 s and 0.98% at 24.49 s were measured from the pixel-wise changes displayed in the maps. Across five stimulation blocks, unsmoothed regional ΔCBV values of 6.57 ± 0.55% contralaterally and −0.15 ± 0.39% ipsilaterally were obtained (mean ± s.e.m. across five trials in one mouse; Fig. 3I, J).

Across mice, ΔCBV values of 6.52 ± 0.78% contralaterally and 0.55 ± 0.26% ipsilaterally were measured in the plane selected for the largest five-trial average contralateral response in each animal (mean ± s.e.m.; n = 6 mice; Fig. 3K). A significant contralateral increase and between-hemisphere difference were detected (both adjusted P = 0.00110), whereas a significant ipsilateral increase was not detected (adjusted P = 0.0837; two-sided paired t-tests with Holm correction). Because the same responses were used for selection and testing, this comparison was treated as exploratory; an all-plane sensitivity analysis is reported in Methods. Access through the native scalp and skull was demonstrated by vascular imaging across W1–W8 and functional mapping at W7. Preparative demands were reduced by avoiding cranial surgery, through which repeated measurements and studies of vascular development could be facilitated.

### Agent-assisted exploration of vascular dynamics

Agent assistance was then extended from imaging configuration and functional mapping to iterative exploration of vascular dynamics (Supplementary Video 5; Fig. 4A). Spectral analysis was used to examine response features beyond location and mean amplitude. Separate awake, cranial-window fUS recordings were used, with stimulus-associated maps and traces (Fig. S4A–E, J), repeated-block amplitudes (Fig. S4F) and first-peak latencies (Fig. S4G, H). At the researcher’s request, spectra and band-power summaries were computed, relevant literature was retrieved and spatial maps were generated by the agent. The outputs were reviewed and each subsequent analysis was directed by researchers.

**Fig. 4.**
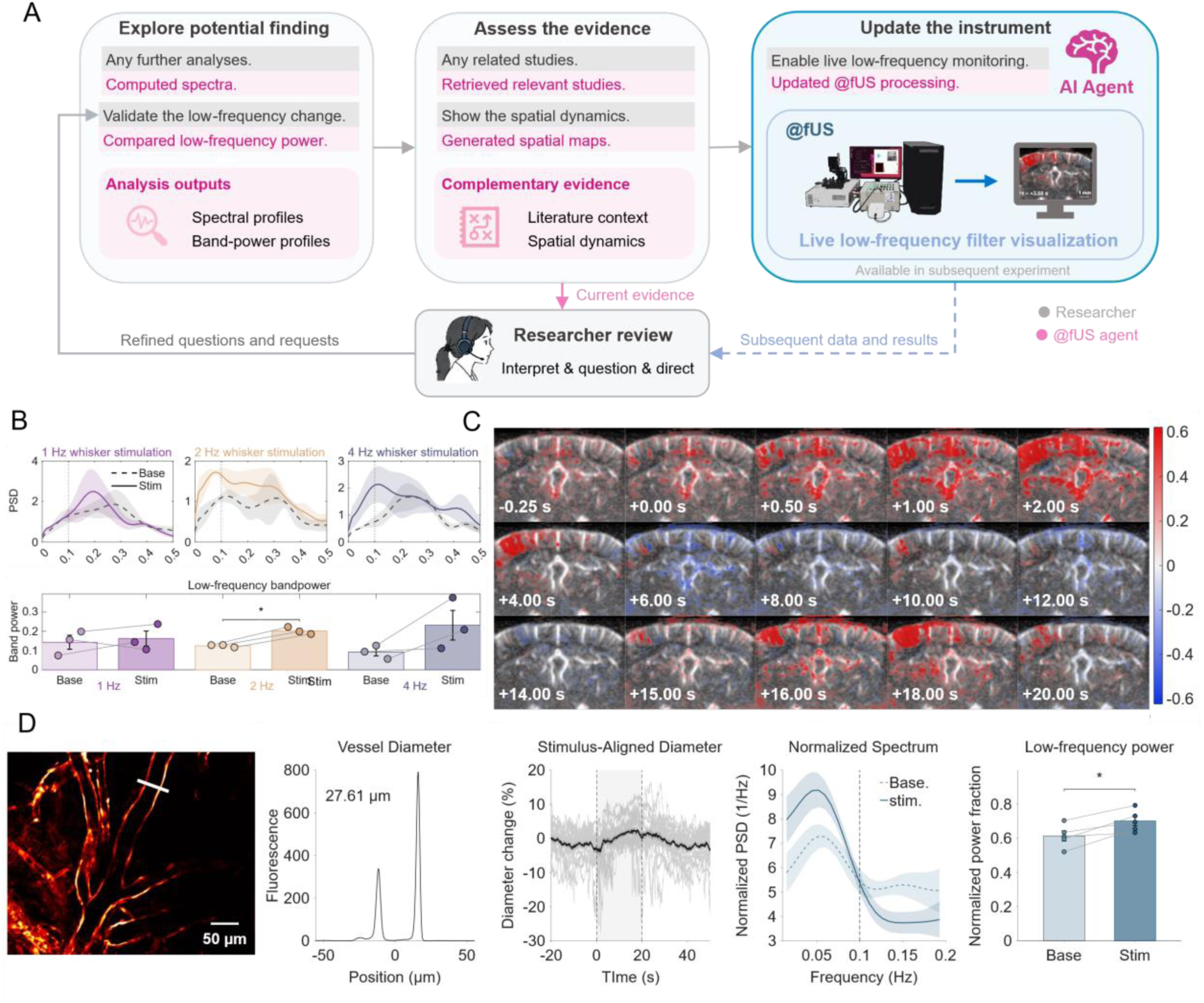
Agent-assisted exploration identifies low-frequency vascular changes and informs processing updates. A, Researcher-directed exploratory analysis, evidence assessment and incorporation of low-frequency visualization into @fUS for subsequent experiments. Researcher requests and review are denoted by grey; agent actions are denoted by pink. B, Baseline (dashed) and stimulation (solid) fUS power spectral density (PSD) estimates, and paired band-power measurements over 0.02–0.15 Hz during 1-, 2- and 4-Hz whisker stimulation (n = 3 mice; five trials per condition averaged within each mouse). Mean ± s.e.m. is shown by curves, bars and shading; individual mice are represented by connected points. Two-sided paired t-tests within each frequency, without correction across frequencies: P = 0.645, 0.0112 and 0.173, respectively. C, signed 0.02–0.15-Hz components of continuous fUS signals around stimulation onset, obtained by offline, non-causal filtering. Positive and negative components are indicated by red and blue. Filtered signals are displayed in these maps; band power, statistical activation, physiological onset and propagation delays are not established by this visualization. D, Complementary two-photon measurements: endothelial fluorescence, a transverse intensity profile for diameter estimation, stimulus-aligned diameter changes, normalized spectra and paired low-frequency power fractions. Twenty-seven vessel segments from six recordings in five mice were included. Mouse–trial averages are shown by thin grey traces; their mean ± s.e.m. is shown by the black trace and shading. Spectra were calculated using paired 20-s baseline and stimulation epochs, with frequency interpolation and normalization over 0.02–0.20 Hz. Power fractions were calculated as power over 0.01–0.125 Hz divided by power over 0.01–0.20 Hz. Between-mouse mean ± s.e.m. is shown by spectral curves, shading and bars; individual mice are represented by connected points. Two-sided paired t-test, P = 0.0153. *P < 0.05. Independent spectral information was not increased by interpolation. Separate awake, head-fixed cranial-window preparations were used for the two modalities, as specified in Methods. Scale bars, 1 mm (C) and 50 µm (D).

An increase in fUS power over 0.02–0.15 Hz was detected during 2-Hz stimulation, from 0.1243 at baseline to 0.2024 during stimulation (P = 0.0112; n = 3 mice; Fig. 4B). Significant baseline– stimulation differences were not detected at 1 Hz (0.1430 versus 0.1624; P = 0.645) or 4 Hz (0.0923 versus 0.2314; P = 0.173). Comparisons were performed using two-sided paired t-tests on mouse-level averages of five trials, without correction across the three frequencies. The 2-Hz result was treated as an exploratory observation for further examination; differences between stimulation frequencies were not tested. To examine the spatial distribution of slow signal changes, the signed 0.02–0.15-Hz component of continuous fUS recordings was mapped by the agent (Fig. 4C). In the displayed sequence, early changes were distributed broadly across the imaged brain and were not confined to SSp-bf. Spatially varying positive and negative components were observed over time.

On the basis of literature retrieved by the agent, slow stimulus-evoked changes in vessel caliber were considered as a candidate contribution to the low-frequency fUS response^42,43^. Vessel-diameter dynamics were examined directly using complementary two-photon recordings (27 vessel segments from six recordings in five awake mice; Fig. 4D). A gradual increase in mean diameter during stimulation and subsequent recovery were observed in stimulus-aligned traces obtained by tracking endothelial fluorescence boundaries. In paired 20-s baseline and stimulation epochs, an increase in the low-frequency power fraction was observed in all five mice, from 0.614 ± 0.030 to 0.702 ± 0.028 (mean ± s.e.m.; two-sided paired t-test, P = 0.0153). This fraction was defined as power over 0.01–0.125 Hz divided by power over 0.01–0.20 Hz. The two-photon observations were considered consistent with a contribution of slow vascular responses to low-frequency signal changes.

Following researcher review, low-frequency visualization was incorporated into the @fUS processing pipeline for subsequent experiments (Fig. 4A). The configuration-and-feedback workflow in Fig. 2A was thereby extended: an analytical observation was linked to a candidate explanation, complementary vascular measurements and an additional processing function. Parameter comparison, functional analysis and exploration of findings were thus supported by agent assistance, and the processing capabilities available for subsequent measurements were informed by experimental evidence.

## Discussion

@fUS demonstrates how a scientific instrument can integrate physical measurement with the practical knowledge required to design, execute and interpret experiments. Its central contribution is the co-design of high-performance, open imaging hardware and an agent layer with executable skills and persistent structured memory. Together with transcranial imaging, these components address technical, preparative and operational barriers to the wider adoption of fUS. Configurations, procedures and experimental history become resources that can be inspected, reused and revised within a researcher-directed workflow, helping researchers translate scientific objectives into experiments and use their outcomes to guide further investigation^9,44^.

The hardware provides the measurement foundation for this integration. @fUS combines 128-channel acquisition, 16-bit RF digitization at 125 MHz, sustained transfer of 16 GB/s to host memory and GPU-based reconstruction. These capabilities extend those of mini-fUS which uses 64 channels and 12-bit digitization at 50 MHz^26^. The nominal digitization resolution and sampling rate exceed those of the original Verasonics Vantage and us4R, and match Vantage NXT, for which the manufacturer specifies sustained host transfer of up to 6.6 GB/s^45^. @fUS combines this acquisition specifications with an open architecture built for fUS, exposing hardware configuration, RF data and reconstruction choices within the same experimental framework. Integrated commercial systems such as Iconeus One and programmable research platforms have advanced access to fUS^23^; @fUS adds direct coordination of these capabilities through its agent layer. The improved vascular visibility and contrast in superficial and deep brain regions (Fig. 2) support the practical value of the integrated design, while retained RF data permit observations to be re-examined through alternative reconstructions.

Accessibility also depends on the biological preparation^23,46–50^. @fUS detected cerebral vessels and stimulus-associated hemodynamic responses through the native scalp and skull after hair removal alone (Fig. 3), reducing the need for cranial surgery in the mouse preparations examined. Previous studies established transcranial fUS in rodents by using contrast agent or EDTA^51–54^; here, this capability is incorporated into an open platform with configurable acquisition, reconstruction and agent-assisted operation^23^. The anatomical localization and stimulus-associated timing of the responses support the recovery of functional information through the native preparation. By reducing surgical preparation, this approach could facilitate repeated measurements and broaden participation in longitudinal studies, provided that performance is established for the relevant ages and experimental conditions.

Agent-compatible design is a natural fit for fUS because acquisition, image formation and analysis depend on interrelated computational choices. Executable skills connect scientific objectives to documented instrument and analytical functions, while persistent memory preserves the context needed across successive interactions. Text and voice interfaces allow researchers to inspect proposed operations and direct their execution with less manual workflow assembly. SmartTrap and EIMS illustrate the value of feedback-controlled optical manipulation and adaptive microscopy, respectively^1,39^. Agent access across the acoustic measurement pipeline is enabled in @fUS by a hardware architecture designed for agent integration, with programmable interfaces for electronics-level control, raw RF data, beamforming, clutter filtering and functional analysis. An unexpected observation can therefore be examined at different stages of image formation and used to guide subsequent measurements. Embedding this coordination within the instrument simplifies interaction while preserving the methodological choices available to the researcher.

The low-frequency analysis illustrates how this architecture can support an investigation emerging from the data (Fig. 4). Agent-assisted spectral analysis identified a stimulation-associated change and provided a basis for considering whether slow changes in vessel caliber contribute to the fUS response^42,43,55^. Complementary two-photon recordings showed a slow mean diameter response and an increased low-frequency power fraction, consistent with this candidate explanation. However, the 20-s optical epochs limited spectral resolution, and frequency interpolation did not add independent spectral information^56^. The evoked response itself may contribute to the low-frequency component; the analyses did not isolate additional oscillatory activity or establish the neural origin or causal significance of the spectral changes^57^. Because the two modalities measured different vascular signals and frequency intervals, their findings provide complementary evidence rather than a direct replication of the same metric. The example connects analytical exploration with a question for further testing, while the processing update makes the resulting visualization available for subsequent experiments.

Several limitations define the next stage of development. The study establishes agent–instrument integration, while dedicated adaptation of language and vision models to ultrasound experimentation remains an area for further development. Available data span a limited range of probes, preparations, image appearances and instrument states, and the skill library mainly supports foundational acquisition and common analyses. Broader use will require additional skills and systematic evaluation across users and laboratories, including task completion, parameter-selection errors and recovery from unfamiliar conditions. Interpretations and proposed operations should remain linked to inspectable evidence and validated tools^5,35^. The present findings establish researcher-directed experimentation; fully autonomous investigation will require further validation of experimental planning, physical execution and the handling of unexpected outcomes.

Open release provides a route to expand both the capabilities and deployment of @fUS. Laboratories could contribute annotated recordings, acquisition configurations and executable procedures, supported by versioning, provenance, compatibility information and documented validation. Successful workflows and recorded failures could both inform improvements, allowing practical expertise to accumulate alongside data and code^58^. By lowering technical and operational barriers, @fUS could become a widely deployable tool across neuroscience laboratories, substantially increasing the volume and diversity of functional imaging data. Such deployment could also generate datasets linking experimental objectives, instrument settings, analytical decisions and measurement outcomes across diverse biological preparations and experimental conditions. These resources could support the training and evaluation of agents that plan experiments, assess measurement quality and adapt their actions to new conditions, providing a data foundation for future autonomous experimentation. Further integration with robotic positioning, stimulation and complementary recording modalities could support progression towards fully automated cycles of experimental design, execution and analysis within researcher-defined objectives and approved experimental constraints². More broadly, @fUS offers a model for scientific instruments in the AI era, in which wider access to measurement capabilities enables the accumulation of data and practical knowledge needed for increasingly autonomous scientific investigation.

## Methods

### System hardware and assembly

The @fUS system comprised a 128-channel transmit–receive board, a probe connector board and a high-throughput transfer board installed in a workstation (Fig. 1 and Fig. S1A). The transmit– receive board integrated ultrasound excitation and 16-bit echo acquisition at 125 MHz, with a single field-programmable gate array (FPGA) coordinating acquisition timing and interfaces. An STM32 microcontroller managed power and auxiliary board functions. The board operated from a regulated 12-V supply and provided four groups of SMA trigger input/output interfaces, an FPGA mezzanine card (FMC) interface and four optical transceivers with an aggregate link bandwidth of 400 Gbit/s. The workstation used an AMD 9975WX CPU and an NVIDIA L20 GPU and ran Ubuntu 22.04.

The probe connector board supported DLM5 or DLP interfaces. Compatibility was tested with commercial Vermon L22-14vX and L11-5v probes, a custom 15-MHz and 25-MHz probes. The imaging experiments reported here used a custom linear array described below. The connector board connected to the transmit–receive board through six signal connectors (2170903-1, TE Connectivity) and three power connectors (5646955-1, TE Connectivity). A programmable DC supply (QPX600DP, Aim-TTi, or DP832A, RIGOL) provided positive and negative transmit-voltage rails.

The transfer board received data over the optical links and transferred it to host memory through two PCIe Gen4 ×8 interfaces, with a combined theoretical peak bandwidth of approximately 32 GB/s. The fUS-optimized streaming path sustained a measured transfer rate of 16 GB/s to host memory. High-bandwidth memory (HBM) buffered interruptions associated with PCIe DMA interrupts while preserving continuous streaming. Because RF data were streamed during acquisition, an entire power Doppler ensemble did not need to reside in front-end memory; ensemble length could instead be configured subject to downstream memory and processing capacity. The probe, transmit–receive board and workstation were connected through the electrical and optical interfaces described above.

### Instrument control and reconstruction

The host application exposed acquisition and reconstruction through human-readable TXT and CSV configuration files (Fig. 2A and Table S1). Hardware settings included transmit amplitude, element-specific delays, receive settings, digital filters and firmware-exposed register values. Reconstruction settings included angle selection, the beamforming grid, coherent compounding, SVD clutter filtering, power Doppler ensemble length and display scaling. Configurations could be loaded without recompiling the host software or FPGA firmware and were retained with the corresponding data.

The host application parsed each configuration and exchanged commands and status information with the acquisition hardware through a bidirectional interface. RF packets were transferred by PCIe direct memory access into a host circular buffer and decoded using one or more CPU threads. Payload dimensions, packet and frame lengths, and sequence continuity were checked before reconstruction; invalid frames were flagged. Frame headers retained sequence numbers, hardware-trigger states and system-status information.

Validated RF frames were staged in pinned host memory and transferred asynchronously to the GPU for digital filtering, delay-and-sum beamforming, coherent angular compounding, SVD clutter filtering and power Doppler estimation. Reception, decoding, transfer and reconstruction were pipelined. The application displayed power Doppler images and acquisition status and supported storage of RF data, reconstructed images and configurations for offline analysis.

### Agent control and analysis

A locally deployed agent framework accessed @fUS through wrappers around the host software and analysis tools (Fig. S1B, C). Natural-language requests, including transcribed voice input, were converted into plans. Approved operations were executed through Model Context Protocol (MCP)-compatible tools and task-specific skills. Instrument control and analysis ran in the local @fUS environment. Knowledge resources and experimental data were stored locally, and the framework also supported cloud-based services. The standalone application remained available for manual operation. The framework was adapted to work with ChatGPT 5.6, ChatGPT 6 and Kimi.

The agent used a planner to decompose requests, a tool router to select functions and a validation layer to check commands and configuration files with the host parser. In the workflow shown in Fig. 2A, documented skills linked the requested imaging task and laboratory constraints to proposed acquisition and reconstruction settings. Scalar software settings were represented in TXT files and hardware-register tables in CSV files. Available tools supported configuration editing, status inspection, acquisition control, live image retrieval, reconstruction changes, parameter sweeps, image-quality comparison, atlas matching, analysis and reporting. Proposed operations and parameter changes required researcher approval before execution or configuration loading; researchers could approve, revise or reject them. Images and analytical outputs were available for review before further operations were requested.

The agent’s built-in knowledge retrieval and memory functions accessed locally stored system documentation, protocols, example configurations, representative datasets and selected literature. Persistent memory retained previous configurations, experimental history and user preferences, which could be overridden by new instructions. Reusable skills specified task objectives, inputs, tools, procedures and outputs. Skills covered system setup, acquisition, reconstruction, image inspection, parameter comparison, atlas-guided positioning, experimental planning, analysis and reporting.

For image optimization, alternative SVD rank-rejection fractions and display-scaling settings were generated through predefined sweeps. A configuration was selected by the researcher after inspecting the outputs (Fig. 2A and Fig. S1C). For the angular comparison in Fig. 2B–E, independently acquired recordings were analyzed by the agent. For the complementary comparison in Fig. S3, the same recording was used across retrospective angular reconstructions within each mouse. All fUS data analysis and statistical testing, including the supporting analyses, were performed by the agent in MATLAB via matlab-mcp-server or the MATLAB Engine API for Python, under researcher approval. For Fig. 3, these analyses included vascular image quantification, ensemble-length comparison, trigger alignment, regional response extraction, GLM mapping and mouse-level statistical summaries. Analytical definitions and inclusion rules were those specified in the corresponding Methods subsections. Two-photon analysis and statistical testing were performed separately. Shared-memory access supported analysis during acquisition.

In the exploratory workflow in Fig. 4A, researcher requests directed spectral estimation, baseline–stimulation band-power comparisons, literature retrieval and visualization of slow spatial signal changes. Complementary two-photon measurements were assessed alongside the fUS results. Following review, low-frequency visualization was incorporated into the processing pipeline for subsequent experiments. The retrospective offline maps in Fig. 4C were analyzed separately from this later implementation.

### Animals and experimental preparations

Male C57BL/6J mice, including all neonatal age groups, were used for fUS experiments. Transgenic mice expressing mCherry in vascular endothelial cells were used for two-photon imaging. The main experiments comprised cranial-window vascular imaging (Fig. 2), intact-scalp and intact-skull vascular and functional imaging (Fig. 3), and separate awake cranial-window fUS and two-photon measurements (Fig. 4). Procedures were approved by the Committee for the Use of Laboratory Animals at the Guangdong Institute of Intelligence Science and Technology, China, and followed institutional guidelines. Ages and sample sizes are reported in the corresponding figure legends and analysis.

For Fig. 2 and the fUS experiments in Fig. 4, a cranial window was implanted under intraperitoneal Zoletil (50 mg/kg) and xylazine (2.6 mg/kg) anesthesia, administered at 10 µl/g body weight. The skull over the imaging region was removed with the dura intact and replaced with a poly(4-methyl-1-pentene) (TPX) window. A custom head plate was attached for fixation. Mice received postoperative anti-inflammatory treatment and analgesia and recovered for 1 week. Vascular imaging in Fig. 2 was performed under 1% isoflurane delivered in oxygen at 0.4 L/min (RWD Life Science). Whisker-stimulation recordings in Fig. 4 were obtained in awake, head-fixed mice23.

Transcranial imaging in Fig. 3 was performed through the intact scalp and skull. Vascular images were acquired under 1% isoflurane delivered in oxygen at 0.4 L/min, as in Fig. 2. Functional recordings during whisker stimulation were obtained in 7-week-old mice under intraperitoneal Zoletil–xylazine anesthesia, administered at 3.5 µl/g body weight. Hair was shaved and residual hair removed with depilatory cream only in mice older than 2 weeks; mice aged 2 weeks and younger required no hair removal. Mice younger than 2 weeks were stabilized in a custom holder. Recordings used for the retrospective comparison in Fig. S3 were obtained under the same Zoletil– xylazine regimen as the functional experiments in Fig. 3.

For two-photon imaging, a glass coverslip and head plate were implanted under Zoletil–xylazine anesthesia, with the dura preserved. After recovery, mice were imaged awake and head-fixed during whisker stimulation (Fig. 4).

### Ultrasound acquisition and angular compounding

Imaging used a custom linear array with a 110-µm pitch and a nominal center frequency of 15 MHz; transmissions used the same frequency. For 21-angle acquisition, each cycle comprised 21 plane-wave transmissions from −10° to +10° in 1° increments at a pulse repetition frequency of 11.44 kHz. This yielded an acquisition-cycle rate of approximately 544.8 Hz. RF channel data were streamed for beamforming and coherent compounding.

For the angular-compounding comparison in Fig. 2B–E, RF recordings were acquired independently for each configuration: 21 angles from −10° to +10° in 1° increments, 11 angles over the same range in 2° increments, or seven angles at −6°, −4°, −2°, 0°, +2°, +4° and +6°. Images acquired at the configured transmit angles were coherently compounded within each acquisition cycle. Power Doppler images used 300 compounded frames and removal of the leading 20% of SVD components by rank. The SVD rank-rejection fraction denotes the proportion of components removed after ordering singular values from largest to smallest; it is not an amplitude threshold. Seven mice contributed at least 100 power Doppler images per mouse and condition.

The SVD comparison in Fig. 2F used the same 21-angle data with rank-rejection fractions of 15%, 25% and 40%, holding other reconstruction settings constant. The transcranial vascular images and age-group comparisons in Fig. 3B, C used removal of the leading 20% of SVD components by rank27. The comparison in Fig. 3D used fractions of 10%, 25% and 40%. Functional recordings in Figs. 3 and 4 used all 21 angles. In Fig. 3F, the same RF recordings were reconstructed with non-overlapping ensembles of 50 or 300 compounded frames, corresponding to integration durations of 0.0917238 or 0.5503428 s from the recorded timing and output rates of 10.9 or 1.82 Hz, respectively. Other functional recordings used 50-frame ensembles.

### Whisker stimulation and synchronization

Whiskers were stimulated with a motor controlled by a Teensy 4.1 microcontroller. Host software set the excursion range and movement speed; the range was adjusted to engage most whiskers, and cyclic stimulation was delivered at 1, 2 or 4 Hz. Transcranial functional recordings in Fig. 3 used 2-Hz stimulation; the awake fUS recordings in Fig. 4 used 1-, 2- and 4-Hz stimulation. For both fUS and two-photon imaging, recordings began with a 50-s baseline followed by five blocks of 20-s stimulation and 30-s recovery. After the fifth 30-s recovery period, recording continued for an additional 30–60 s. A digital output remained high throughout each stimulation epoch. For fUS, this signal was connected by coaxial cable to the transmit–receive board and recorded in the acquisition headers.

### Vascular image analysis

Visible vessel area fraction, labelled vessel density or vessel area in the figures, was defined as the percentage of an analysis region of interest (ROI) occupied by Doppler-visible vascular pixels (Figs. 2E and 3C). Segmentation combined multiscale vessel enhancement (0.08–0.50 mm), Otsu thresholding (multiplier, 0.9; minimum threshold, 0.08), an intensity cutoff of 0.1 and morphological cleanup59. A common segmentation threshold was used within each comparison. This image-derived fraction was used to quantify vascular visibility.

For angular comparisons (Fig. 2E), matched analysis and background ROIs were used across conditions. Visible vessel area fraction and contrast were measured separately in the upper and lower halves of the analysis ROI. Apparent vessel diameter was estimated from the −3-dB width of manually selected vascular profiles, with the median valid width calculated for each frame. Vessel-to-background contrast was calculated as 10*log10(Pv/Pb), where Pv and Pb were mean powers within the vascular and background ROIs. Frame-level measurements were averaged within each mouse and condition before statistical testing.

For the transcranial cross-plane analysis, five vascular reference images were obtained by averaging the 10-s baseline preceding each stimulation. Vessel enhancement was performed on an isotropic grid, normalized across slices and mapped back to the native grid for area measurements. Visible vessel area fraction was averaged across the five baseline images. Segmentation sensitivity was evaluated by changing the primary threshold by ±20%.

### Spatial and temporal image characterization

For the complementary comparison in Fig. S3, the same 21-angle RF recording from each mouse was retrospectively reconstructed using 7, 11 or 21 angles. This comparison was used to reduce confounding by changes in physiological state between independently acquired recordings. For each reconstruction, temporal coefficient-of-variation maps were calculated as 100 × stdₜ[P(t)]/meanₜ[P(t)], where P(t) is the pixel-wise power Doppler signal. Inter-frame difference and spatial-correlation time courses summarized variation within the analyzed vessel mask. The representative sequence summarized in Results comprised 190 frames and 189 adjacent-frame pairs over 98.28 s, with a recorded frame interval of 0.55 s. Adjacent-frame measurements were averaged within this sequence; successive frames were not treated as independent biological replicates.

To assess whether differences in temporal intensity variation were accompanied by changes in vessel localization, transverse sections along representative cortical and deep-brain vessels were used to examine signal time courses and position variability (Fig. S3I–Q). The position-jitter value reported in Results refers to the central sampled section of the cortical vessel. The section labelled 0 in the figure served as the reference for peak cross-correlations and associated delays; whole-vessel spectra described signal-frequency content. These analyses assessed reconstruction-dependent spatial consistency and temporal variability.

### Transcranial functional responses

For the functional recordings in Fig. 3E–K, power Doppler images were assigned ensemble-center timestamps and aligned to hardware-recorded stimulation onsets. Trigger states were sampled at 0.0917238-s intervals, corresponding to 50 compounded frames. For 300-frame ensembles (0.5503428 s), an ensemble was classified as stimulation if any trigger sample within its interval was high. The ensemble-length comparison in Fig. 3F used the fifth trial from mouse 1, plane 1, with a common valid primary somatosensory barrel field (SSp-bf) mask and without temporal smoothing or nuisance correction. For each reconstruction, ΔCBV was averaged over the stimulation period within this single trial, and baseline temporal standard deviation was calculated on the native sampling grid. These summaries were not averages across repeated trials. This comparison characterized sampling and baseline variability in one trial; it was not an equivalence test of functional detection sensitivity.

Bilateral SSp-bf ROIs were defined by translation and scaling to the Allen Mouse Brain Common Coordinate Framework, version 3 (CCFv3), and classified as ipsilateral or contralateral to stimulation60. Regional ΔCBV was estimated from spatially averaged linear power as 100 × [P(t) − P₀]/P₀, where P₀ was mean power during each trial’s 10-s pre-stimulus baseline. Five individually normalized trials were aligned and averaged (Fig. 3J). Continuous traces used the baseline before the first stimulation (Fig. 3I). An approximately 0.5-s trailing moving average was used only for display; response amplitudes were calculated from unsmoothed signals across the full anatomical ROI (Fig. 3K and the all-plane sensitivity analysis). Slice selection and mouse-level aggregation are specified under Statistical analysis and reproducibility.

### Activation mapping

For Fig. 3G, H, pixel-wise general linear models (GLMs) were fitted to log-power changes referenced to median baseline log-power pooled across non-stimulation samples in the 30-s pre-stimulus periods47. The design matrix contained a stimulus regressor convolved with a single-gamma hemodynamic response function (delay, 1.0 s; time constant, 0.7 s; order, 3), its temporal derivative, an intercept, linear and quadratic drift terms, and low-frequency cosine terms with a cutoff of 0.008 Hz.

A global nuisance time course was estimated as the spatial median of task-and-drift residuals standardized by their median absolute deviation. This time course and its temporal derivative were orthogonalized against the task and drift design before joint fitting. Slice-level first-order autoregressive pre-whitening accounted for temporal autocorrelation. Positive task effects were tested with one-sided t-tests and Benjamini–Hochberg false discovery rate correction within each slice (q < 0.05), and displayed as statistical Z maps.

Dynamic maps in Fig. 3H showed five-trial-averaged ΔCBV after removal of task-protected global nuisance components from log-power signals and conversion to relative power. Signals were normalized to each trial’s 10-s baseline. Images at 0.25-s target intervals were selected from the nearest acquired time points without interpolation. Only positive changes within the full-session GLM significance mask were displayed; the mask was not re-estimated at each time point. Maps used a common 0–33% display range. Accompanying ROI means were calculated over the entire anatomical SSp-bf from numerical ΔCBV values. Fig. 3G shows GLM-derived statistical Z maps at ten atlas-derived anterior–posterior positions from −1.70 to +0.10 mm relative to bregma, with 200-µm spacing. Anatomical SSp-bf masks were defined separately at each plane, allowing ROI area to vary with cross-sectional anatomy. Activated SSp-bf fraction was defined as positive significant area divided by the full anatomical ROI area. Regional ΔCBV was calculated across the entire anatomical ROI, without restricting it to significant pixels. The all-plane analysis included 9, 9, 10, 9, 10 and 9 valid contralateral SSp-bf planes in mice 1–6, respectively (56 total); absent ROIs were treated as missing. After the prior quality exclusions for mouse 3, planes were not excluded according to response amplitude. Cross-plane summaries were averaged within each mouse before calculation of the six-mouse mean and s.e.m..

### Extended response characterization

Fig. S4 summarizes whisker-evoked responses at 1, 2 and 4 Hz using response maps, continuous and stimulus-aligned traces, block-wise amplitudes and peak-fraction latencies. Repeated-block amplitudes were expressed relative to the first block. Latencies were measured relative to stimulation onset at 20%, 50%, 80% and 100% of the first response peak; values beyond the 3-s plotting range were marked at the upper boundary. Raw and smoothed traces were displayed separately. These time-domain summaries were treated separately from the fUS spectral measure in Fig. 4B and the optical power fraction in Fig. 4D.

### Exploratory fUS spectral analysis

The exploratory analysis in Fig. 4B used awake cranial-window fUS recordings during 1-, 2- and 4-Hz whisker stimulation in three mice, with five trials per condition. Baseline and stimulation spectra were compared, and power over 0.02–0.15 Hz was summarized across trials within each mouse before paired testing. This band-power measure was analyzed separately from the two-photon power fraction defined below. For Fig. 4C, continuous pixel-wise fUS signals were filtered non-causally over 0.02–0.15 Hz and aligned to stimulation.

### Two-photon imaging and vessel diameter analysis

For the complementary measurements in Fig. 4D, two-photon OIR recordings were imported into MATLAB with Bio-Formats at a frame interval of 65.779 ms. A reference image was generated by two-dimensional translation registration and averaging of 25 frames acquired 20–40 s after recording onset. Vessel ROIs and longitudinal axes were defined manually. Transverse fluorescence profiles were averaged along each vessel, smoothed with a Gaussian kernel and used to locate the two wall peaks with subpixel precision. Their separation was converted to micrometers using the image calibration.

Frame-wise translation was corrected before diameter extraction. Failed registrations and invalid diameter measurements were excluded. Internal gaps were linearly interpolated without a maximum gap length. A valid pair comprised the same vessel and trial during the 20-s baseline (−20 to 0 s) and stimulation (0–20 s) epochs; each epoch required at least 80% valid measurements before interpolation and only finite values after interpolation. Included vessels had at least four valid pairs, at least 80% valid measurements across the recording, a registration failure rate of no more than 20% and acceptable reference-image quality. The review interface displayed reference images and vessel cross-sections and did not mask animal identity or experimental condition. Diameter changes were expressed relative to the median valid diameter before the first stimulation, and trials were aligned to trigger rising edges. The dataset comprised 27 vessel segments from six recordings in five mice. For response plots, vessels were averaged within recordings, then recordings from the same mouse were averaged by trial number to obtain mouse–trial traces.

Two-photon spectral analysis used paired 20-s pre-stimulation (−20 to 0 s) and stimulation (0– 20 s) epochs. Signals were linearly detrended and Hann-windowed before one-sided periodogram estimation. Spectra were interpolated in frequency for band integration and display. Displayed spectra were normalized by total power over 0.02–0.20 Hz and plotted over 0.01–0.20 Hz. The separately reported low-frequency power fraction was power over 0.01–0.125 Hz divided by power over 0.01–0.20 Hz. Spectra and power fractions were averaged across trials, vessels and recordings to obtain one summary per mouse.

### Statistical analysis and reproducibility

Mice were the independent biological replicates; frames, pixels, trials, vessels and slices were within-animal measurements. Angular-compounding comparisons in Fig. 2E used two-sided paired t-tests on mouse-level summaries (n = 7), with Holm correction across the three angle pairs for each region and metric. Representative temporal values from Fig. S3 and the ensemble comparison in Fig. 3F were descriptive. For Fig. 3K (n = 6 mice), one plane per mouse was selected by the largest contralateral SSp-bf stimulation-period ΔCBV averaged across five trials. Each hemisphere was represented by its five-trial mean from that plane. Two-sided paired t-tests compared baseline with stimulation in each hemisphere and stimulus-induced changes between hemispheres, with Holm correction across these three comparisons. Baseline-normalized comparisons were equivalent to tests against zero. Because selection and testing used the same responses, this analysis was exploratory.

A sensitivity analysis averaged all eligible SSp-bf planes equally within each mouse before applying the same three paired tests. Mean stimulation-period ΔCBV was 4.30 ± 0.61% contralaterally and 0.50 ± 0.19% ipsilaterally; Holm-adjusted P values were 0.00208 for the contralateral increase and between-hemisphere difference, and 0.0457 for the ipsilateral increase. Exact two-sided sign-flip tests enumerated all 64 sign assignments for each six-mouse comparison. After Holm correction, exact P values were 0.09375 for all three contrasts in both the response-selected and all-plane analyses. Thus, significance depended on the test used in this small cohort. Cross-plane summaries used mouse-level averages and did not test differences between anterior– posterior positions.

Exploratory fUS band-power comparisons in Fig. 4B used two-sided paired t-tests on mouse-level trial averages (n = 3), comparing baseline with stimulation separately at each stimulation frequency, without correction across the three frequencies. No between-frequency test was used to support the spectral interpretation. Two-photon power-fraction comparisons in Fig. 4D used two-sided paired t-tests on mouse-level summaries (n = 5). The fUS and two-photon measurements were analyzed separately because their signals, frequency bands and preparations differed.

For Figs. 2–4, group summaries are mean ± s.e.m. unless stated otherwise. Age-group vessel area fractions in Fig. 3C show between-mouse standard deviation (three mice per group across W1– W8). Representative fUS response curves in Fig. 3J show s.e.m. across five trials. Two-photon diameter-response curves show s.e.m. across mouse–trial traces, whereas spectral shading and power-fraction error bars show between-mouse s.e.m. Paired points in the main quantitative comparisons represent mice. Significance thresholds were *P < 0.05, **P < 0.01, ***P < 0.001 and ****P < 0.0001, using adjusted P values where specified.

## Data availability

The data supporting this study is available at https://doi.org/10.5281/zenodo.22907752.

## Code availability

The code supporting this study is available at https://github.com/HQArrayLab/agentic_tfUS.

## Acknowledgments

This work is supported in part by the National Key Research and Development Program of China (Grant No. 2023YFC2410900), the National Natural Science Foundation of China (32371151, 32571281, and 92570202), Guangdong High Level Innovation Research Institute (2021B0909050004), the Hong Kong Research Grants Council Senior Research Fellowship (SRFS2526-5S03), Collaborative Research Fund (C5053-22 GF), General Research Fund (15126524 and 15224323), National Key Research and Development Program of Ministry of Science and Technology of China (2023YFC2410900), Research Center for Non-invasive Brain Computer Interface (1-CE0M), and Research Institute of Smart Ageing (1-CDJM and 1-CDMZ), Zhejiang Provincial Natural Science Foundation of China (LQK26C090002).

## Author contributions

Conceptualization: Z.H.Q., Z.H.C., N.L.; @fUS design and engineering: Z.H.C., N.L.; Agent development: Z.Q.L., Z.X.Y., Y.M.W., Z.H.C.; Experiments: Z.H.C., N.L.; System test and workflow optimization: Z.H.C., Z.R.L., C.Y.F., L.S., W.L., H.Z.L., H.Y.Y., T.Y.W, B.X.L., Q.X.X., Y.Z., Y.L., H.S.W., W.J.B., X.D.L., Y.C., X.Z., B.L.; Data analysis: Z.H.C., L.N.; Manuscript writing: Z.H.Q., Z.H.C., N.L., L.S., Z.Q.L., Z.X.Y., Y.M.W., Z.R.L., C.Y.F., L.S., W.L., H.Z.L., H.Y.Y., T.Y.W, B.X.L., Q.X.X., Y.Z., Y.L., H.S.W., W.J.B., X.D.L., Y.C., X.Z., B.L.; Funding acquisition: Z.H.Q., Y.M.W., L.S., W.J.B.; Supervision: Z.H.Q., Y.M.W., L.S.

## Supporting figures and table

**Fig. S1.**
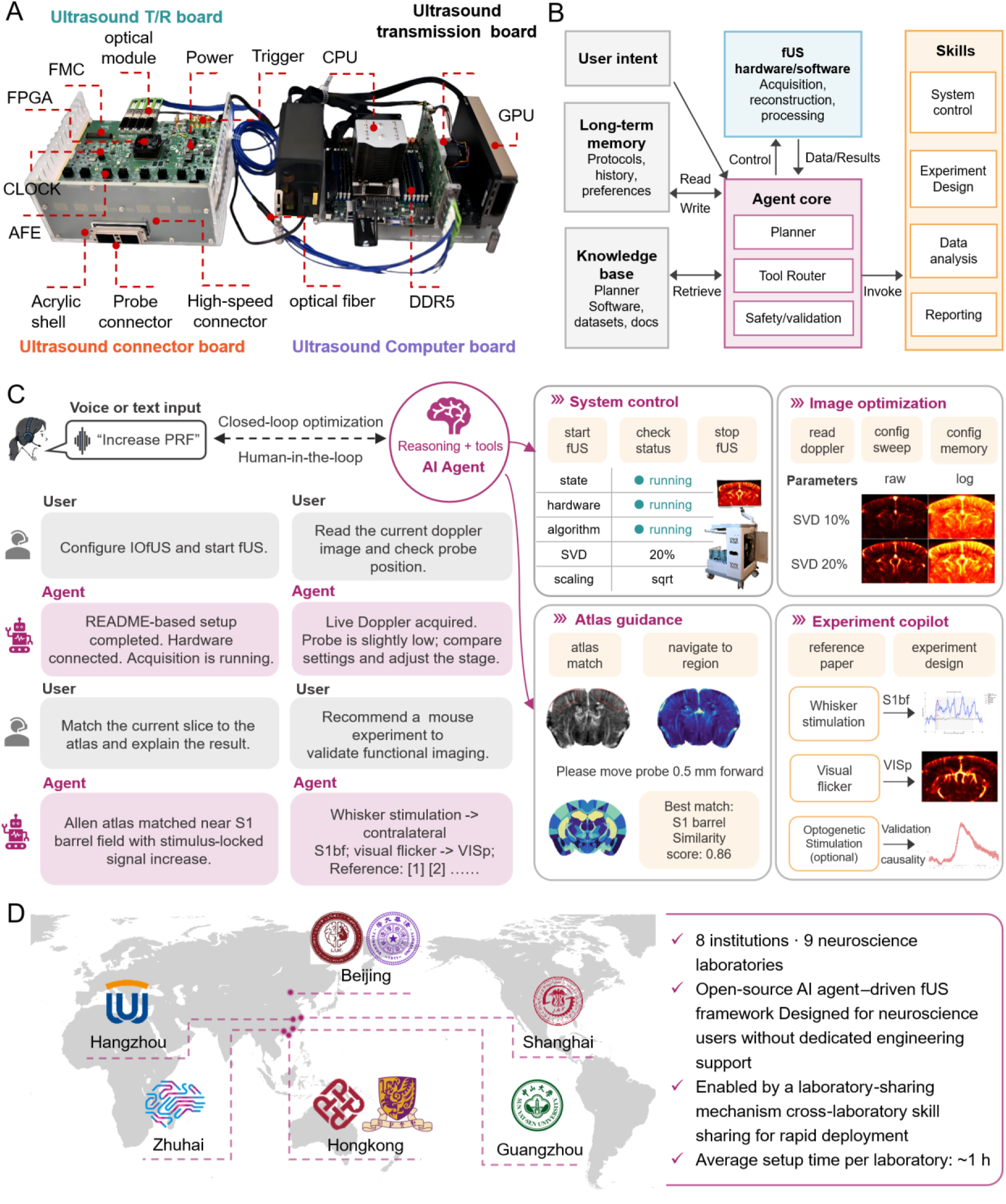
Hardware and agent implementation for researcher-supervised experiments. A, Ultrasound transmit–receive and probe-connector electronics linked to the transfer board and workstation, expanding Fig. 1. B, Agent architecture connecting user intent, instrument tools, stored knowledge and persistent experimental context through planning, tool routing and validation. Instrument control, experimental design, analysis and reporting are supported by executable skills. C, Illustrative text and voice interactions and tool views for system control, image inspection, SVD-rank comparison, atlas guidance and experimental planning. Researcher approval is required for proposed operations. The workflow is illustrated by dialogue and interface values. D, Reported deployment network of nine neuroscience laboratories at eight institutions. Deployment scope is described in this panel; controlled user-performance evaluation was not performed. Hardware and agent implementation are described in Methods. AFE, analog front end; FPGA, field-programmable gate array; FMC, FPGA mezzanine card; CPU, central processing unit; GPU, graphics processing unit; SVD, singular value decomposition; T/R, transmit–receive.

**Fig. S2.**
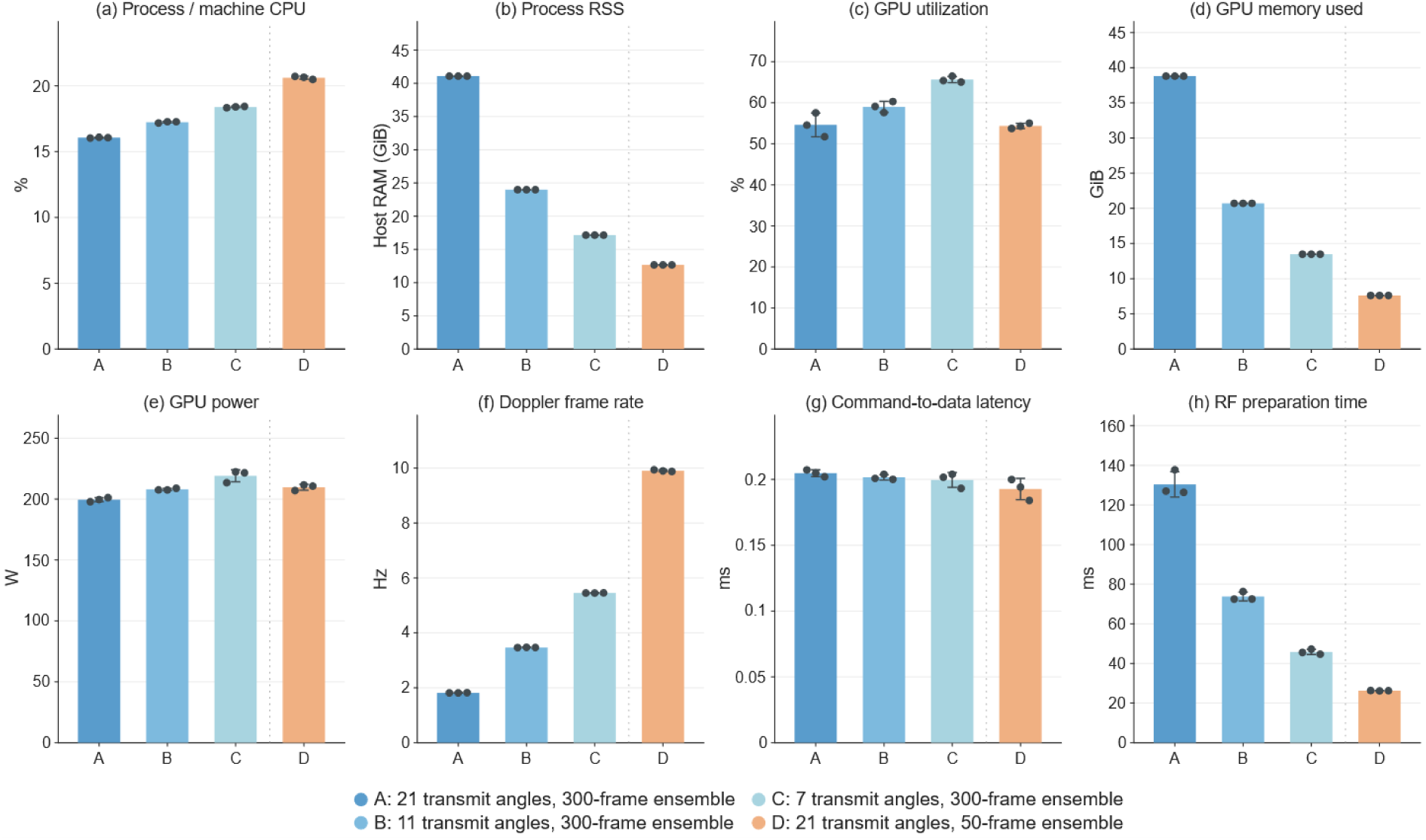
System resource use and imaging performance under different parameter settings. Independent resource tests were performed at a fixed pulse repetition frequency. Configurations A–D on the horizontal axes were defined as 21 angles with 300-frame ensembles, 11 angles with 300-frame ensembles, 7 angles with 300-frame ensembles and 21 angles with 50-frame ensembles, respectively. A, Receiving-process CPU usage as a percentage of total online logical-core capacity. B, Host RAM occupied by the receiving process, measured as resident set size (RSS). C–E, GPU utilization (C), memory use (D) and power (E), measured for the entire GPU. F, Real-time Doppler imaging frame rate, measured from the Qt image-update counter. The measured frame rate was affected by Qt memory reads; the raw-data acquisition frame rate was unaffected. G, Latency from the logged user request to start acquisition to the first recorded data reception. H, Cumulative mean RF preparation time. Mean ± sample standard deviation across three independent runs per configuration is shown by bars and error bars; individual runs are indicated by points. Resource metrics and imaging frame rates were summarized over the valid 30 to 160-s interval after acquisition was initiated by the user. One first-reception latency and one cumulative RF preparation mean through approximately 160 s, including startup, were obtained per run.

**Fig. S3.**
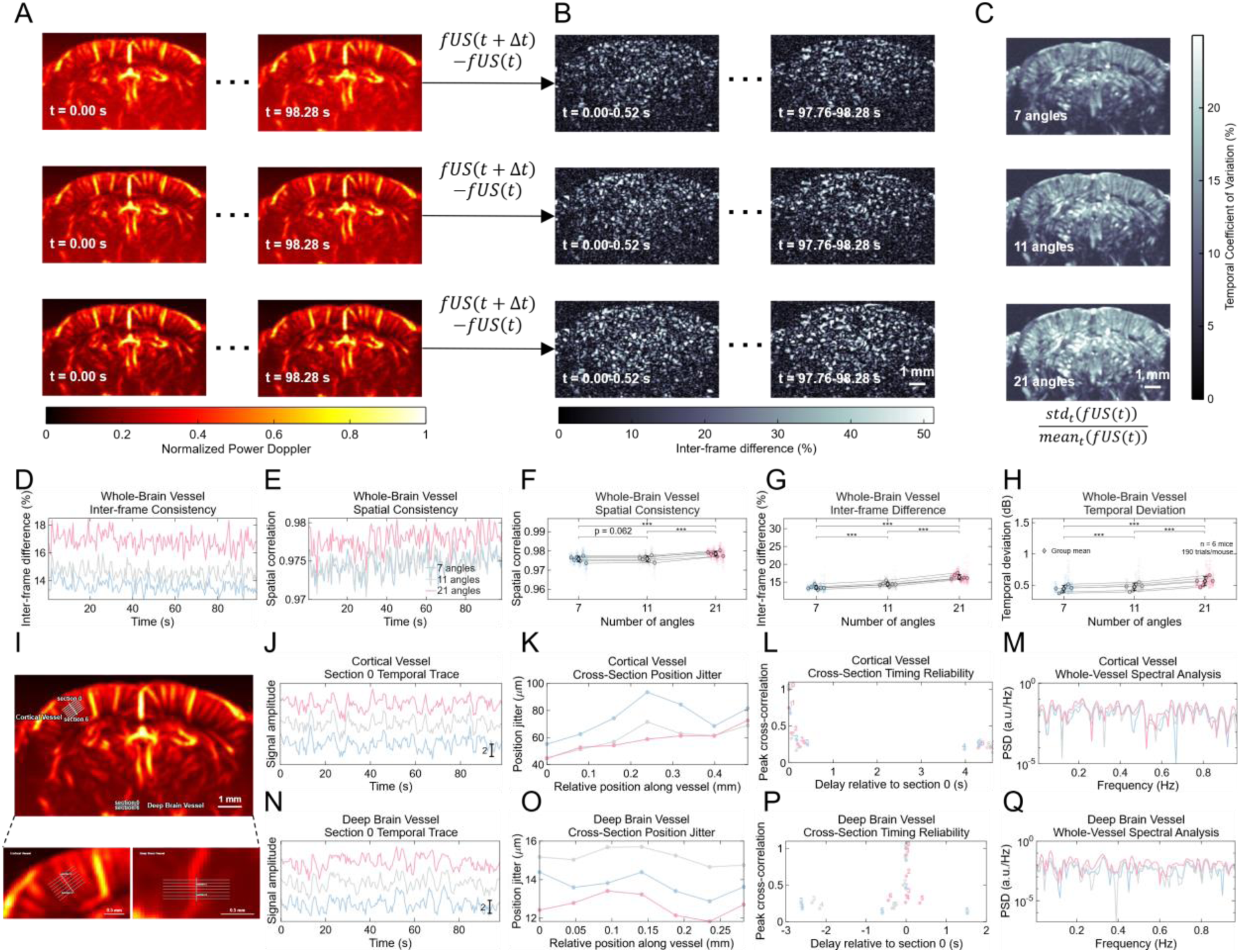
Spatial consistency and temporal variability across angular reconstructions. Complementary retrospective comparison of temporal intensity variation and spatial localization across angular configurations. To reduce confounding by changes in mouse physiological state, the same 21-angle RF recordings were reconstructed using 7, 11 or 21 angles. Recordings were acquired under intraperitoneal Zoletil–xylazine anesthesia (3.5 µl/g body weight), as used for the functional experiments in Fig. 3 (Methods). A–C, Power Doppler sequences (A), inter-frame difference maps (B) and temporal coefficient-of-variation maps (C) for seven-, eleven- and twenty-one-angle reconstructions. Temporal coefficients of variation are standard deviations divided by means, expressed as percentages. D, E, Inter-frame differences (D) and spatial correlations (E) over time within the analyzed vessel mask. F–H, Spatial correlation (F), inter-frame difference (G) and temporal deviation (H) across reconstruction configurations. Paired measurements from six mice are indicated by linked summaries. The representative 190-frame recording in A–E is described by the numerical sequence-level summaries in Results, which are distinguished from these mouse-level summaries. I, Cortical and deep-brain vessels with transverse sampling sections indicated in the enlargements. J–M, Cortical-vessel signals at reference section 0 (J), cross-section position variation along the vessel (K; the central section is quantified in Results), peak cross-correlations and associated delays relative to section 0 (L), and whole-vessel power spectra (M). N–Q, Corresponding reference-section signals (N), position variation (O), cross-correlations and delays (P), and spectra (Q) for the deep-brain vessel. The visibility and contrast metrics in Fig. 2 are complemented by these measures, through which reconstruction-dependent variability is characterized without an independent temporal ground truth. Analysis is described in Spatial and temporal image characterization in Methods. Seven, eleven and twenty-one angles are denoted by blue, grey and pink. Times are relative to recording onset. Scale bars, 1 mm (B, C and overview in I) and 0.5 mm (enlargements in I).

**Fig. S4.**
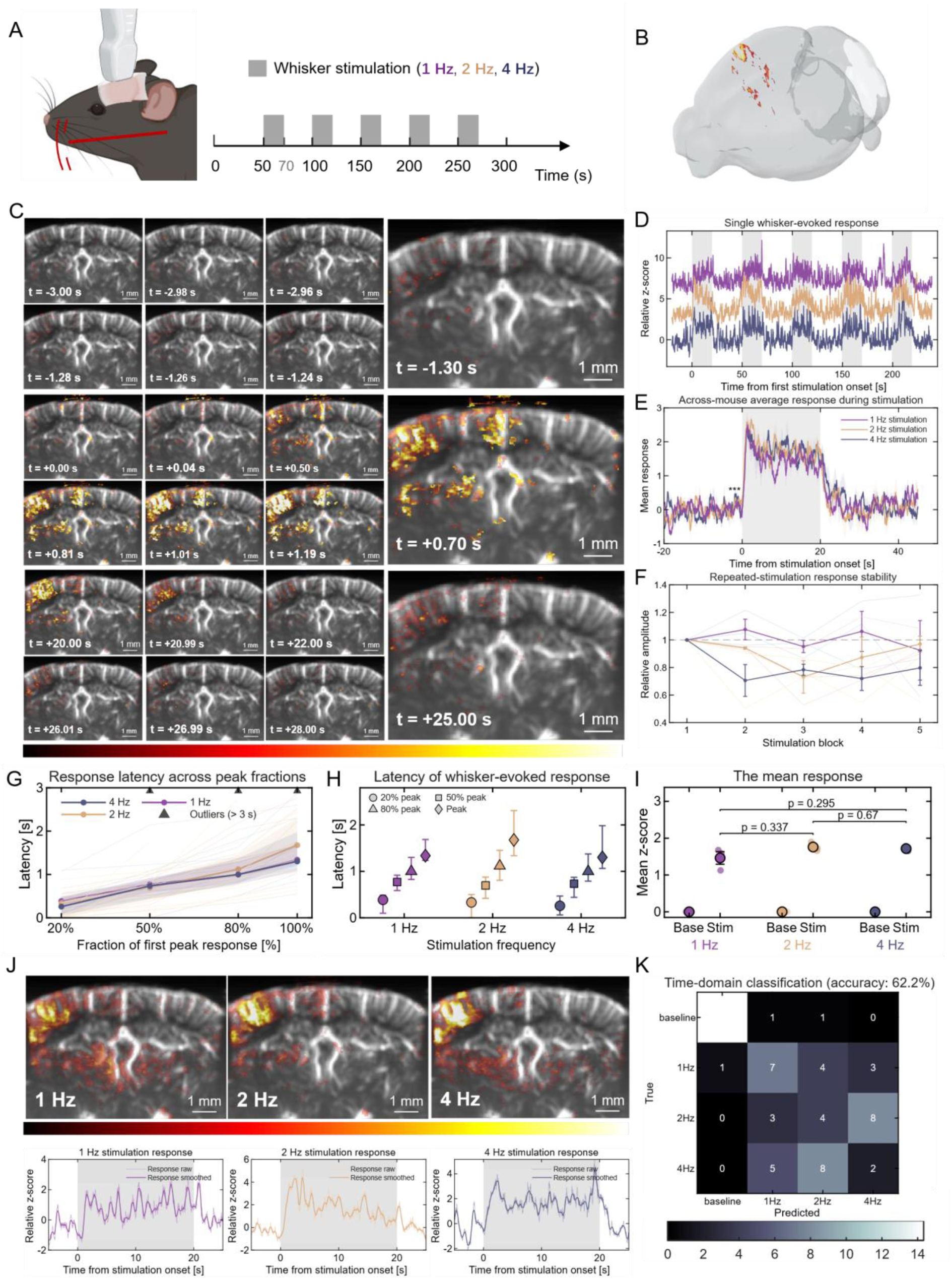
Time-domain characterization of whisker-evoked fUS responses. Time-domain characterization of the awake cranial-window fUS recordings used for the spectral exploration in Fig. 4 (n = 3 mice; five trials per stimulation frequency). These recordings were separate from the anesthetized transcranial cohort in Fig. 3. A, Whisker stimulation at 1, 2 or 4 Hz, comprising a 50-s baseline and five blocks of 20-s stimulation and 30-s recovery. B, Three-dimensional localization of the displayed response. C, Representative response overlays before, during and after stimulation; selected frames are enlarged at right. Times are relative to stimulation onset. D, E, Representative responses across stimulation blocks (D) and across-mouse mean stimulus-aligned responses (E). Stimulation is indicated by grey shading. F, Response amplitudes across five blocks, normalized to the first block. G, H, Latencies to 20%, 50%, 80% and 100% of the first response peak, plotted against peak fraction (G) and grouped by stimulation frequency (H). Values beyond the 3-s display range are indicated by upper-boundary markers in G. I, Baseline and stimulation response summaries at each frequency. J, Response maps and corresponding raw and smoothed traces at each frequency. K, Exploratory time-domain classification, with true labels in rows and predicted labels in columns. Response characterization is described in Methods; neither the frequency contrasts nor the classifier is used to support the low-frequency interpretation in Fig. 4. Stimulation at 1, 2 and 4 Hz is denoted by purple, orange and blue. Scale bars, 1 mm (C, J).

**Table S1.** Instrument interfaces relevant to agent integration. Qualitative comparison supporting the architecture in Fig. 1 and the configuration workflow in Fig. 2A. Documented platform capabilities are distinguished from integration work for an external agent. Public product and software documentation is summarized in the Iconeus One and Verasonics descriptions^45^; The implementation described in Methods is summarized for @fUS. Absence of a capability cannot be established from missing documentation for a specific external-agent interface. Interfaces are compared; imaging quality, agent performance and deployment time were not assessed in a controlled benchmark.

| Instrument capability | Iconeus: Application-oriented fUS system | Verasonics: General-purpose programmable ultrasound platform | @fUS: Structured openness for agent integration |
| --- | --- | --- | --- |
| **Perception: Machine-readable data and state** | Provides real-time fUS images and integrated analysis results. Public documentation of interfaces for external agents to access raw data and low-level operating states is limited. | Provides access to raw RF and processed data, but users must package device states, error information and quality metrics into agent-readable interfaces. | Provides unified access to raw RF, IQ, Power Doppler, trigger information, frame numbers, packet-loss information, buffer states and system operating states, allowing the agent to observe both experimental results and instrument status. |
| **Computation: Computation-ready real-time data pathway** | Includes an optimized fUS processing pipeline, but interfaces allowing external agents to access the internal real-time computational pipeline are not publicly documented. | Supports custom online processing and GPU computation, but users must implement the complete fUS processing workflow and its agent interfaces. | Streams 16-bit data acquired at 125 MHz to the GPU through high-speed optical links, PCIe and shared memory. The pipeline integrates beamforming, SVD, Power Doppler and machine-readable quality-control outputs. |
| **Control: Broad, structured and bounded instrument control** | Parameters are adjusted primarily through integrated software and a graphical user interface, providing stable operation. However, the range of actions callable by external agents is limited or not publicly documented. | Provides extensive control over transmission, reception and processing through MATLAB or C, but the agent must generate or invoke complex programs. | Scalar parameters are defined in TXT files, while large angle, delay and channel tables are defined in CSV files. The agent modifies configurations through constrained tools without changing application code, the graphical interface or FPGA firmware. |
| **Robustness: Verified and recoverable operation** | Internal system integration reduces operational errors, but interfaces for external state readback and software-based recovery are not publicly documented. | Users can implement custom control, but parameter validation, execution confirmation and fault-recovery logic require additional development. | Configurations are validated before loading, and registers and lookup tables are read back after execution. When an abnormal state occurs, the system supports software abort, soft reset and restoration of the last-known-good configuration. |
| **Context: Continuity and provenance** | Data and analysis results are retained within the vendor's workflow, but interfaces allowing external agents to retrieve and maintain the complete experimental context are not publicly documented. | MATLAB/C states, scripts and data can be saved, but cross-session experimental associations, agent memory and operational provenance must be implemented by the user. | Shared memory exposes current acquisition, computation and quality-control states. File-based interfaces maintain links among experimental configurations, data indices, analysis results, agent actions and human approvals, allowing experiments to be resumed, inspected and continued. |

## References

1. Selin, M. et al. SmartTrap: automated precision experiments with optical tweezers. Nat Methods 23, 1368–1378 (2026).

2. Boiko, D. A., MacKnight, R., Kline, B. & Gomes, G. Autonomous chemical research with large language models. Nature 624, 570–578 (2023).

3. Vriza, A., Prince, M. H., Zhou, T., Chan, H. & Cherukara, M. J. Operating advanced scientific instruments with AI agents that learn on the job. npj Comput Mater 12, 160 (2026).

4. Chen, Z., et al. An agentic artificially intelligent X-ray scientist. Nat Mach Intell 8, 1075– 1086 (2026).

5. M. Bran, A., et al. Augmenting large language models with chemistry tools. Nat Mach Intell 6, 525–535 (2024).

6. Szymanski, N. J. et al. An autonomous laboratory for the accelerated synthesis of inorganic materials. Nature 624, 86–91 (2023).

7. Burger, B. et al. A mobile robotic chemist. Nature 583, 237–241 (2020).

8. Kusne, A. G. et al. On-the-fly closed-loop materials discovery via Bayesian active learning. Nat Commun 11, 5966 (2020).

9. Sim, M. et al. ChemOS 2.0: An orchestration architecture for chemical self-driving laboratories. Matter 7, 2959–2977 (2024).

10. Pinkard, H. et al. Pycro-Manager: open-source software for customized and reproducible microscope control. Nat Methods 18, 226–228 (2021).

11. Steiner, S. et al. Organic synthesis in a modular robotic system driven by a chemical programming language. Science 363, eaav2211 (2019).

12. Macé, E. et al. Functional ultrasound imaging of the brain. Nature methods 8, 662–664 (2011).

13. Nunez-Elizalde, A. O. et al. Neural correlates of blood flow measured by ultrasound. Neuron 110, 1631–1640.e4 (2022).

14. Landemard, A., Krumin, M., Harris, K. D. & Carandini, M. Brainwide blood volume reflects opposing neural populations. Nature 1–9 (2026) doi:10.1038/s41586-026-10350-9.

15. Rabut, C. et al. 4D functional ultrasound imaging of whole-brain activity in rodents. Nat Methods 16, 994–997 (2019).

16. Brunner, C. et al. A Platform for Brain-wide Volumetric Functional Ultrasound Imaging and Analysis of Circuit Dynamics in Awake Mice. Neuron 108, 861–875.e7 (2020).

17. Boido, D. et al. Mesoscopic and microscopic imaging of sensory responses in the same animal. Nat Commun 10, 1110 (2019).

18. Li, B. et al. Circuit mechanism for suppression of frontal cortical ignition during NREM sleep. Cell 186, 5739–5750.e17 (2023).

19. Griggs, W. S. et al. Functional ultrasound neuroimaging reveals mesoscopic organization of saccades in the lateral intraparietal area. Nat Commun 16, 8752 (2025).

20. Baranger, J. et al. Bedside functional monitoring of the dynamic brain connectivity in human neonates. Nature communications 12, 1080 (2021).

21. Zhu, J. et al. Functional Imaging of Developing Brain in Mice and Non-Human Primates. bioRxiv 2024–06 (2024).

22. Faure, F. et al. Cot-side functional imaging in neonates for early neurodevelopment monitoring using functional ultrasound (fUS) connectivity imaging and the combination of fUS with diffuse optical tomography (fUS-DOT): A feasibility study. Developmental Cognitive Neuroscience 77, 101652 (2026).

23. Brunner, C. et al. Whole-brain functional ultrasound imaging in awake head-fixed mice. Nature Protocols 16, 3547–3571 (2021).

24. Brown, M. D. et al. Four-dimensional computational ultrasound imaging of brain hemodynamics. Science Advances 10, eadk7957 (2024).

25. Huang, Y.-A. et al. OfUSA: OpenfUS Analyzer, a versatile open-source framework for the analysis and visualization of functional ultrasound imaging data across animal models. 2025.09.16.676515 Preprint at 10.1101/2025.09.16.676515 (2025).

26. Yang, J. et al. Miniaturized and accessible functional ultrasound imaging system for freely moving mice. 2025.04.24.650351 Preprint at 10.1101/2025.04.24.650351 (2025).

27. Demené, C. et al. Spatiotemporal Clutter Filtering of Ultrafast Ultrasound Data Highly Increases Doppler and fUltrasound Sensitivity. IEEE Transactions on Medical Imaging 34, 2271– 2285 (2015).

28. Le Meur-Diebolt, S. et al. Robust functional ultrasound imaging in the awake and behaving brain: A systematic framework for motion artifact removal. Imaging Neuroscience 4, IMAG.a.1191 (2026).

29. Lambert, T., Brunner, C., Montaldo, G. & Urban, A. PyfUS: Python-based open-source software for the analysis of functional ultrasound imaging data. Neurocomputing 651, 130899 (2025).

30. Hayashi, S. et al. brainlife.io: a decentralized and open-source cloud platform to support neuroscience research. Nat Methods 21, 809–813 (2024).

31. Zhang, G. et al. SVD-SN2N: Self-Inspired Noise2Noise Learning for Denoising Log-Compressed SVD-Filtered Ultrasound Imaging. 2026.09.10.750668 Preprint at 10.64898/2026.09.10.750668 (2026).

32. Işık, E. B. et al. Grand challenges in bioinformatics education and training. Nat Biotechnol 41, 1171–1174 (2023).

33. Oellermann, M. et al. Open Hardware in Science: The Benefits of Open Electronics. Integr Comp Biol 62, 1061–1075 (2022).

34. Wenzel, T. Open hardware: From DIY trend to global transformation in access to laboratory equipment. PLOS Biology 21, e3001931 (2023).

35. Miao, J., Davis, J. R., Zhang, Y., Pritchard, J. K. & Zou, J. Reimagining research papers as interactive and reliable AI agents. Nature https://doi.org/10.1038/s41586-026-11044-y (2026) doi:10.1038/s41586-026-11044-y.

36. Huang, K. et al. Autonomous biomedical research with an artificial intelligence agent. Science 393, eadz4351 (2026).

37. Zhang, H. G., Eckmann, P., Miao, J., Mahon, A. B. & Zou, J. The Virtual Biotech: A multi-agent AI framework for therapeutic discovery and development. Science eaeg6779 (2026) doi:10.1126/science.aeg6779.

38. Lu, C. et al. Towards end-to-end automation of AI research. Nature 651, 914–919 (2026).

39. Gao, H., et al. Embodied Intelligence Enables Agentic Exploration in Microscopy. Preprint at 10.21203/rs.3.rs-8617009/v1 (2026).

40. Marin, Z. et al. Microscope control with a natural language agent. 2026.09.15.751723 Preprint at 10.64898/2026.09.15.751723 (2026).

41. CopilotJ. https://copilotj.chat/#/home.

42. Mayhew, J. E. W. et al. Cerebral Vasomotion: A 0.1-Hz Oscillation in Reflected Light Imaging of Neural Activity. NeuroImage 4, 183–193 (1996).

43. Hill, R. A. et al. Regional Blood Flow in the Normal and Ischemic Brain Is Controlled by Arteriolar Smooth Muscle Cell Contractility and Not by Capillary Pericytes. Neuron 87, 95–110 (2015).

44. Wilkinson, M. D. et al. The FAIR Guiding Principles for scientific data management and stewardship. Sci Data 3, 160018 (2016).

45. Villani, F., et al. Validation of a Software-Defined 100-Gb/s RDMA Streaming Architecture for Ultrafast Optoacoustic and Ultrasound Imaging. Preprint at 10.48550/arXiv.2601.18280 (2026).

46. Urban, A. et al. Real-time imaging of brain activity in freely moving rats using functional ultrasound. Nat Methods 12, 873–878 (2015).

47. Chen, Z. et al. Functional Ultrasound Imaging of Auditory Responses in Comatose Patients. Preprint at 10.1101/2024.12.22.24319283 (2024).

48. Soloukey, S. et al. Functional ultrasound (fUS) during awake brain surgery: the clinical potential of intra-operative functional and vascular brain mapping. Frontiers in neuroscience 13, 1384 (2020).

49. Soloukey, S. et al. Human brain mapping using co-registered fUS, fMRI and ESM during awake brain surgeries: A proof-of-concept study. NeuroImage 283, 120435 (2023).

50. Soloukey, S. et al. Mobile human brain imaging using functional ultrasound. Sci. Adv. 11, eadu9133 (2025).

51. Errico, C. et al. Ultrafast ultrasound localization microscopy for deep super-resolution vascular imaging. Nature 527, 499–502 (2015).

52. Errico, C. et al. Transcranial functional ultrasound imaging of the brain using microbubble-enhanced ultrasensitive Doppler. NeuroImage 124, 752–761 (2016).

53. Tiran, E. et al. Transcranial Functional Ultrasound Imaging in Freely Moving Awake Mice and Anesthetized Young Rats without Contrast Agent. Ultrasound in Medicine & Biology 43, 1679– 1689 (2017).

54. Wang, Z. et al. Acoustic Transparency Enabling Functional Ultrasound Imaging Through Mouse and Human Skulls. 2025.08.22.671878 Preprint at 10.1101/2025.08.22.671878 (2025).

55. Rungta, R. L., Chaigneau, E., Osmanski, B.-F. & Charpak, S. Vascular Compartmentalization of Functional Hyperemia from the Synapse to the Pia. Neuron 99, 362–375.e4 (2018).

56. Mitra, P. P. & Pesaran, B. Analysis of Dynamic Brain Imaging Data. Biophysical Journal 76, 691–708 (1999).

57. Winder, A. T., Echagarruga, C., Zhang, Q. & Drew, P. J. Weak correlations between hemodynamic signals and ongoing neural activity during the resting state. Nat Neurosci 20, 1761– 1769 (2017).

58. Raccuglia, P. et al. Machine-learning-assisted materials discovery using failed experiments. Nature 533, 73–76 (2016).

59. Otsu, N. A Threshold Selection Method from Gray-Level Histograms. *IEEE Transactions on Systems*, Man, and Cybernetics 9, 62–66 (1979).

60. Wang, Q. et al. The Allen Mouse Brain Common Coordinate Framework: A 3D Reference Atlas. Cell 181, 936–953.e20 (2020).

